# A metaproteomics-based meta study of samples from patients with inflammatory bowel disease identifies potential markers for diagnosis and therapy monitoring

**DOI:** 10.1101/2025.11.06.684320

**Authors:** Maximilian Wolf, Luca Knipper, Kay Schallert, Ann-Cathrin Groba, Patrick Hellwig, Corinna Bang, Phillip Rausch, Andre Franke, Dirk Benndorf, Konrad Aden, Alexander Sczyrba, Robert Heyer

## Abstract

Inflammatory bowel disease (IBD) is a chronic intestinal disorder involving recurring inflammation and pronounced microbial dysbiosis. Comprehensive studies with large patient cohorts are required to Identify meaningful biomarker candidates for diagnosing and monitoring IBD. In this large-scale meta-study of over 600 samples based on fecal metaproteomics, our goal was to validate known biomarkers and discover new candidates. We performed bioinformatic reanalysis using the Mascot search engine and MMUPHin for batch effect correction as well as knowledge graph-enhanced data analysis.

We identified 59 protein groups that varied primarily due to disease, rather than laboratory conditions. These included Alpha-1-acid glycoprotein, which was not reported in the original studies. Of these groups, 53 were differentially abundant in at least one of the two validation datasets. Additionally, 23 of the successfully validated protein groups, primarily from human neutrophil vesicles, were found to be significantly associated with remission during treatment in an independent dataset. This finding suggests their potential for disease monitoring.

Validation in other disease contexts, such as non-alcoholic steatohepatitis, diabetes, and colorectal cancer, revealed the necessity of biomarker panels, because individual biomarkers could only distinguish IBD from specific conditions. Our results demonstrate the effectiveness of metaproteomics meta-analyses in discovering and validating biomarker panels and assessing their specificity for IBD.

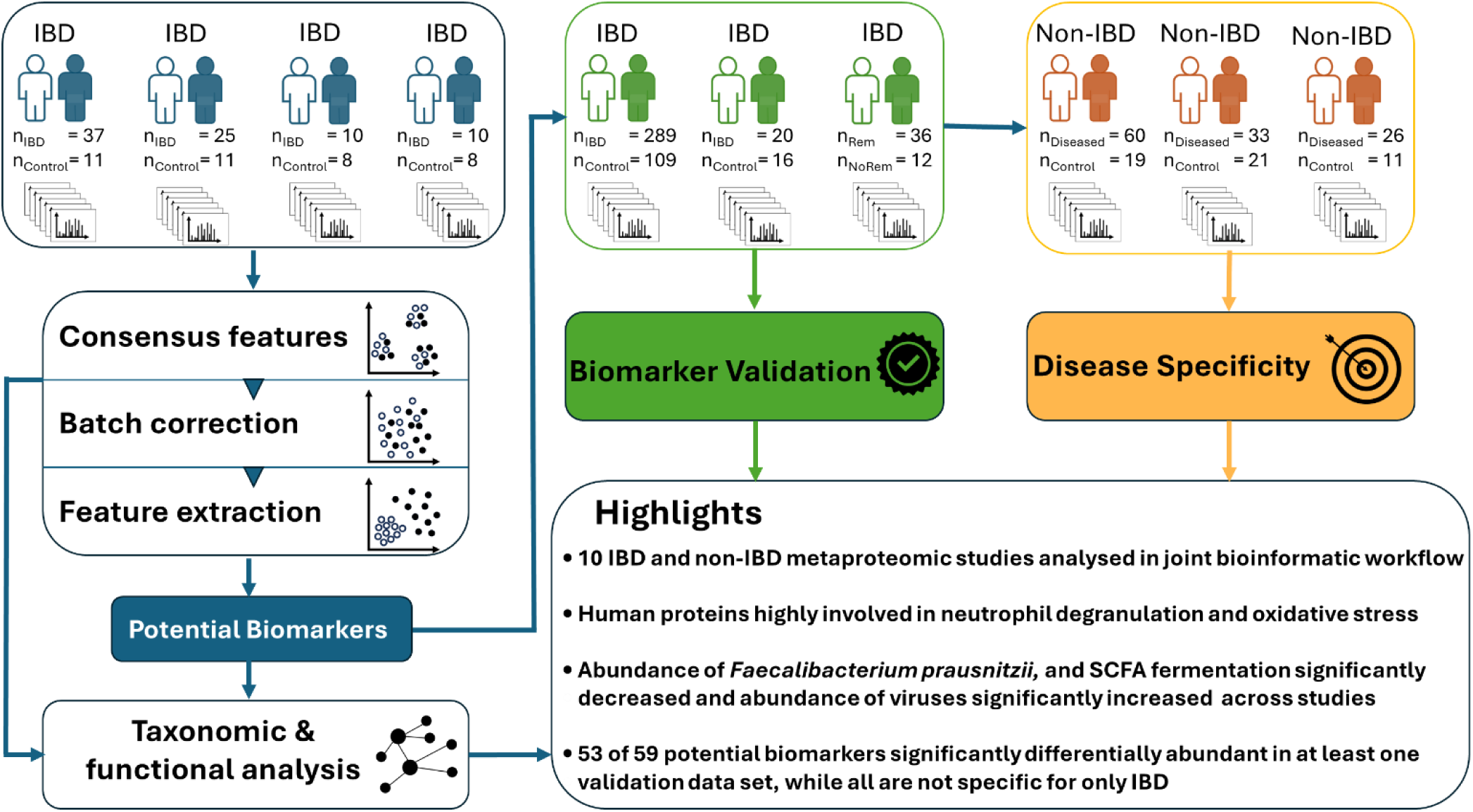

## Introduction

Inflammatory bowel disease (IBD) is a chronic illness of the gastrointestinal tract, which is characterized by a dysregulated immune response leading to a disruption of the gut microbiome and tissue damage, prolonging the inflammatory cascade.

As clinical features like diarrhea or rectal bleeding are nonspecific, diagnosis of IBD is difficult. The gastrointestinal microbiome, as the target of the immune response, has experienced increasing attention for IBD diagnosis and disease monitoring. Several alterations concerning the functional and taxonomic composition of the microbiome of IBD patients were found, using different levels of omics, e.g., metagenomics, metatranscriptomics, and metaproteomics ^1,2^.

Despite growing interest in the role of microbiomes in human health and disease, microbiome studies suffer from several problems:

First, most microbiome studies report associations of a disease with a specific taxon, but often there are inconsistencies in these reported associations. For example, in IBD patients, *Faecalibacterium prausnitzii*, is often reported to be depleted, but an increased abundance of this taxon in IBD has also been described, thereby questioning this association and hampering generalization ^3^. Sources of these inconsistencies could be different wet-laboratory and data analysis workflows. Furthermore, large interindividual differences and small sample size aggravated statistical analysis of associations. Independent validation cohorts, which could be used to verify associations, are rarely used.

Second, the design of microbiome studies often lacks non-healthy control groups, which impedes the evaluation of disease specificity of observed associations. Even though several human biomarkers have been identified, a significant proportion of initial IBD diagnoses are incorrect and instead identified as, e.g., irritable bowel syndrome (IBS), particularly when non-invasive methods are considered ^4^.

To target these two problems, microbiome meta-studies can be conducted. In meta-analyses, raw data from different studies are gathered and analyzed using a joint analysis workflow. Thereby the number of samples is increased and the variation introduced by different data analysis workflows is decreased. Subsequently, microbial associations can be evaluated regarding their consistency and disease specificity.

In their analysis of several metagenomics datasets, Wirbel et al. found that the microbiome of patients with Crohn‘s disease is consistently characterized by a loss of beneficial members of the Clostridiales order ^5^. On the other hand, in a 16S rRNA meta-analysis it was found that the depletion or increased abundance of taxa is mainly not disease specific ^6^. Therefore, the analysis of taxonomical composition might not be sufficient for specific IBD disease diagnosis, leading to the third major problem of microbiome studies: the usage of gene-based methods. 16S rRNA or metagenomics analysis only provide information of the taxonomic composition or functional potential of the microbial community as they assess the genetic material, not the proteins, which exhibit the actual metabolism. This level of information can be assessed with metaproteomics, which has shown its potential and advantages compared to gene-based methods in the context of IBD. Metaproteomics enables the assessment of the hosts immune response by the identification and quantification of human proteins. Compared to metagenomics, metaproteomic profiles demonstrated a stronger association with disease activity in IBD, thereby highliting the advantages of metaproteomics ^7^.

Despite the growing number of clinical metaproteomics studies ^2^, only one meta-analysis of samples from healthy individuals has been conducted to date^8^. This study aimed to identify the healthy metaproteome. The potential of metaproteomics meta-analysis to address the absence of human and microbial biomarkers in IBD has not been explored.

Taken together, there is no study assessing functionally relevant, metaproteomics-derived biomarkers, evaluating and validating their consistency and disease specificity. Therefore, we first analyzed 120 samples from four studies including IBD patients and extracted 53 features (human and microbial) consistently associated with IBD compared to control samples across all four studies. Second, we validated their usefulness as markers for diagnosis and therapy monitoring with three (partially) independent data sets. Finally, we evaluated the specificity of the validated metaproteins in datasets concerning non-IBD diseases, among others, diabetes, Hepatocellular Carcinoma (HCC), or Nonalcoholic Steatohepatitis (NASH).

## Methods

### Study inclusion and data acquisition

We used PubMed to search for studies that published fecal shotgun metaproteomics data of diseased patients and healthy controls. Inclusion criteria into the meta-analysis were, among others use of data-dependent-acquisition, and data publication after 2015. MS data were downloaded from PRIDE ^9^ or MassIVE (http://massive.ucsd.edu). If .mgf files were available these were used, otherwise .raw files were converted into .mgf files using ThermoRawFileParser ^10^. Available patient data, study metadata and repository identifier were collected and integrated (Supplementary File 1).

### Discovery cohorts

Four datasets, including IBD and Controls from the studies of Lehmann et al.( n_IBD_= 25, n_control_= 11) ^11^, Henry et al. (n_IBD_= 10, n_control_= 8) ^12^, Thuy-Boun et al. (n_IBD_= 10, n_control_= 8) ^13^, and Lloyd-Price et al. (n_IBD_= 37, n_control_= 11) ^14^ were selected for multivariate analysis, feature selection, and taxonomical and functional analysis. As the study of Lloyd-Price included multiple measurements of individuals, only one sample of each adult individual was included to preserve statistical independence.

### Validation cohorts

Three distinct datasets were chosen as validation cohorts. First, a hold-out dataset from the study of Lloyd-Price et al. was used (n_IBD_= 289, n_control_= 109). Secondly, an in-house dataset comprising IBD patients and controls residing in the same household was used (n_IBD_ = 20, n_control_= 16). The samples for this dataset were taken from the Inflammatory Bowel Disease Family Cohort (IBD-FC) collected at the University Hospital Schleswig Holstein (Ethical approval number: AZ A117/13). Third, a dataset consisting of IBD patients before and after biologics therapy was used (n_Remission_= 12, n_NoRemission_= 36) ^15^.

### Specificity cohorts

The disease specificity of potential IBD biomarkers extracted from the discovery cohort was evaluated using samples from Sydor et al. (n_HCC_ = 29, n_NASH_ = 31, n_control_ = 19) ^16^, samples from Gavin et al. (n_Diabetes_ = 33, n_Control_ = 21) ^17^ and samples from Lehmann et al. (Gastric Carcinoma (GCA) n_GCA_ = 6, n_IBs_ = 13, Colon Adenoma (CA) n_CA_ = 7).

### Metaproteomic analysis

All .mgf files from discovery, validation and disease specificity analysis were processed using the same bioinformatic analysis. Database searching was done using the Mascot Search engine and a database comprised of a human gut metagenome catalogue and the Uniprot/Swissprot database [January 2019]. For a detailed comparison with other databases for protein identification, please refer to Supplementary Figure 1. The parameters for the mascot searches are specified in Supplementary File 2.

The resulting .dat files were processed using custom java code for filtering, grouping and integration of identified peptides and inferred proteins, respectively. The code is based on the work of Schallert et al.^18^ and publicly available (https://github.com/maximilianwolf1212/MetaproteomicsMetaAnalysis).

Metaproteins were annotated using Prophane (version 6.2.6 ^19^) with the diamond algorithm (e-value: 0.0001, id cut-off: 0.95) against the National Center for Biotechnology Information (NCBI) database (30/03/2022) and the emapper algorithm (seed_ortholog_evalue: 0.001) against the EggNOG database (25/03/2022). Functional metaprotein annotation, e.g., Kyoto Encyclopedia of Genes and Genomes (KEGG) reactions, Enzyme Commission (E.C.) number, Gene Ontology (GO) terms, was attained by a combination of annotations of individual proteins. Taxonomic annotation was assigned by the Lowest Common Ancestor (LCA) rule (LCA threshold: 0.5). The Prophane summary can be found in Supplementary File 3.

Taxonomic and functional annotation of potential biomarkers was additionally done using their identified peptides and the Unipept 4.0 API (version 2.1.3) ^20^. (Supplementary File 4)

### Statistical analysis, visualization and annotation

Multivariate analysis was carried out in Python using pandas, numpy, matplotlib, scipy and skbio, and sklearn. For Principal Coordinates Analysis (PcoA), MinMax scaler was used for preprocessing, followed by computation of a distance matrix based on “Bray-Curtis” distance measure. To quantify the proportional importance of the grouping factors (disease state, study), group differences were assessed using Permonava (999 permutations).

A rank-based ANOVA-like analysis from the SIAMCAT R package was used to quantify the confounding effect of the studies on the metaprotein abundance ^5^. Metaproteins whose variance was strongly confounded by the study (variance explained by study > variance explained by disease) or only minimally explained by disease (<0.2) were excluded from the following analysis and validation.

Protein-protein interaction analysis was carried out using the STRING Application Programming Interface (API) ^21^, by extracting the human protein identifier from the protein groups. Contextualization of human proteins for drug prediction was done using the knowledge graph Pharmebinet ^22^.

For significance testing of the abundance of taxonomic groups, functional pathways, or individual metaproteins across studies, mixed effect models were built using Python’s statsmodels package using the study as random effect.

Significance tests of potential marker metaproteins in individual validation studies were conducted using the Mann-Witney-U test and Benjamini-Hochberg correction, included in the scipy.stats module.

### Batch effect removal

MMUPHin batch effect removal procedure was used to adjust the abundance of metaproteins identified in all discovery cohorts ^23^. MMUPHin uses an extension of the combat algorithm, adjusted to zero-inflated data under consideration of an additional covariate, here disease state.

All analysis scripts are made available as Jupyter Notebooks on github.com/maximilianwolf1212/MetaproteomicsMetaAnalysis.

## Results

### Batch effect correction and features selection improves sample clustering across datasets

In the discovery phase, samples from four different studies were included, resulting in a total of 82 samples from IBD patients and 40 samples from the control group. The samples of the distinct studies were processed with strongly varying experimental workflows (Supplementary File 5). For example, proteins in the study of Lehmann et al. were extracted using phenol, whereas Thuy-Boun et al. used triflic acid. These variations led to a large number of metaproteins identified in only one study (Figure 1A), leading to study-specific metaproteome profiles, which can be observed in a multivariable visualization (Figure 1B). This is supported by the comparison of the PERMANOVA F-statistics, which showed that the largest proportion of variance was attributable to the laboratory of origin rather than disease status (F_disease / F_study = 0.41).

**Figure 1:**
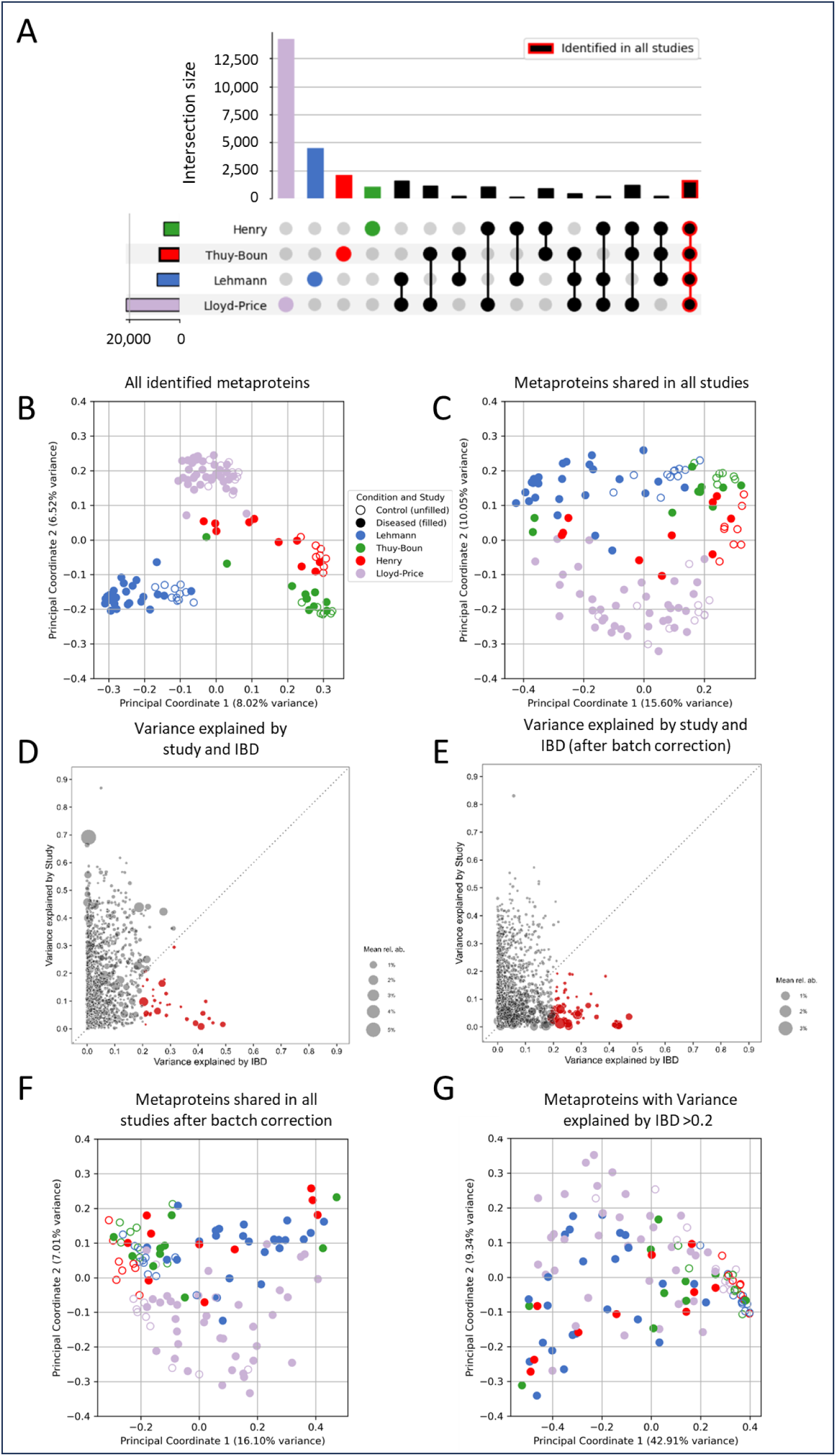
Data integration and feature selection in the discovery cohort of the meta-analysis. A: UpSet plot comparison of sets of metaproteins identified in the different studies. B: PCoA score plot of all metaproteins identified. C: PCoA score plot of metaproteins identified in all studies. D: Variance explained by the study compared to variance explained by IBD for all metaproteins identified in all studies. 32 metaproteins with variance explained by IBD > 0.2 were marked red. E: Variance explained by the study compared to variance explained by IBD for all metaproteins identified in all studies after batch effect correction. 59 metaproteins with variance explained by IBD > 0.2 were marked red. F: PCoA score plot of metaproteins identified in all studies after batch effect correction. G: PCoA score plot of metaproteins all 53 metaproteins with variance explained by IBD > 0.2 after batch effect correction. PcoA – Principal Coordinate Analysis

The separation due to study specific factors was observed in microbiome meta-analyses before. Study-specific microbiome compositions were also detected in meta-analyses of, e.g., 16S-rRNA and metagenomic datasets about colorectal cancer ^24,25^. As strong effects of sample processing, e.g., cell disruption and protein extraction on metaprotein identification from gut samples were observed ^26^, this represent an important source of bias. As the subset of uniquely identified metaproteins declines with the sample size of the respective study, the effect of interindividual differences on the fecal metaproteins becomes clear. Interindividual differences are a key challenge in human microbiome research, as they heavily affect the outcome of medication or dietary interventions ^27^. Meta-analysis can be used to develop microbiome-based stratification strategies for interventional studies by establishing criteria for prediction of highly plastic or highly resistant microbiome structures ^28^, thereby taking into account interindividual differences.

Although “study” was the stronger associated covariate of the metaproteome profile, a separation due to the disease state is recognizable in the samples from the studies of Lehmann et al. and Henry et al., respectively.

Overall, 1,620 metaproteins were identified in all studies (Figure 1A). The summed average relative abundance of these metaproteins is 77.0 ± 9.2%, therefore, they account for a large proportion of all identified spectra. The average abundance of these metaproteins is significantly higher than the abundance of all other sets of shared or uniquely identified metaproteins (p < 0.001) (Supplementary Figure 2). This observation is in line with the results of a metaproteomics ring trial ^29^, which states that the differences between the laboratory workflows used were mostly attributable to low abundant metaproteins. In the metaproteomics meta-analysis of Tanca et al. only seven of 52 identified phyla were identified in all ten datasets analyzed. This finding is a similarly small overlap, and underlines the high proportion of unique features each metaproteomic dataset bears. Additionally, only four phyla were identified in all subjects, underscoring the interindividual differences ^8^.

Even using the set of shared metaproteins, a clear separation due to the different studies can be observed (Figure 1C), however, a separation of samples from healthy and diseased individuals is clearer observable in all studies. The separation of the samples from different studies could be moreover attributed to the enrichment of microbial fractions in at least two of the studies. The largest contributor to the separating dimension is the protein group Chymotrypsin-like Elastase, with a mean abundance of 5.53 ± 6.87% (Supplementary Figure 3), which is the most abundant human protein in stool samples. Enriching the microbial fraction might have lowered the abundance in the samples of Henry et al. and Thuy-Boun et al., whereas the protein was identified in a larger relative abundance in the study of Lehmann et al., in which whole stool samples were used for protein extraction. Consequently, as observable in Figure 1D, the abundance of the greatest part of metaproteins is clearly affected to a large extent by the study-specific effects, and the variance of only a small proportion of the metaproteins is explained by the disease state. Only 32 metaproteins have a proportion of >0.2 of variance explained by IBD.

Batch effect reduction is a commonly used tool in meta-analysis and can possibly enhance study findings ^30^, but could reduce the variance introduced by the disease. Here, we used MUPPHin batch effect removal, which preserves variation between healthy and diseased patients. Reducing the study-specific effects by using MUPPHin, led to an increase from 32 to 59 metaproteins, largely affected by IBD-specific effects (explained variance by IBD > 0.2) rather than study specific effects (Figure 1E). Among these, 31 of 32 from the initial set of metaproteins were present before and after batch effect reduction, showing that this procedure does not reduce biological meaning, but improved the ability for detection of potentially meaningful features. After additional annotation using Unipept, 24 proteins could be identified as human proteins, whereas 35 were considered to be of microbial origin. Using only these 59 proteins, instead of all shared metaproteins after batch effect reduction (Figure 1F, F_disease / F_study = 1.45), a clear separation of samples between healthy and diseased individuals could be observed (Figure 1G). Comparison of PERMANOVA F-statistics underlines the now neglectable importance of the study for the grouping of the patients after batch effect reduction (F_disease / F_study = 11.51). Importantly, microbial proteins are higher abundant in samples from healthy individuals, which are strongly condensed into a small area, whereas the IBD samples are scattered more broadly. As observed with PERMANOVA analysis, the feature selection process led to a substantial decrease in the importance of the technical variation introduced due to the different workflows used in the study and a strong increase in the importance of the disease state (Supplementary Figure 4)

### Hypothesis-driven assessment of human and microbial functions and taxonomies

A central goal of meta-analyses in microbiome research is to refine and prioritize hypotheses by identifying generalizable disease-associated patterns across studies ^31^. Here, we tested a broad set of commonly proposed hypotheses derived from various laboratory approaches, including metaproteomics, metagenomics, RNA sequencing, and microscopy, by analyzing consensus features (metaproteins identified in all discovery studies) within our dataset.

Alterations in taxonomic composition are frequently reported in microbiome studies. In metaproteomic investigations of IBD, an increased abundance of human proteins, particularly those related to immune functions, is commonly observed and has been even correlated with disease severity ^15,32^. Our findings support these reports as the abundance of *Homo sapiens* proteins was significantly elevated in the disease state (p < 0.001, Figure 2A).

**Figure 2.**
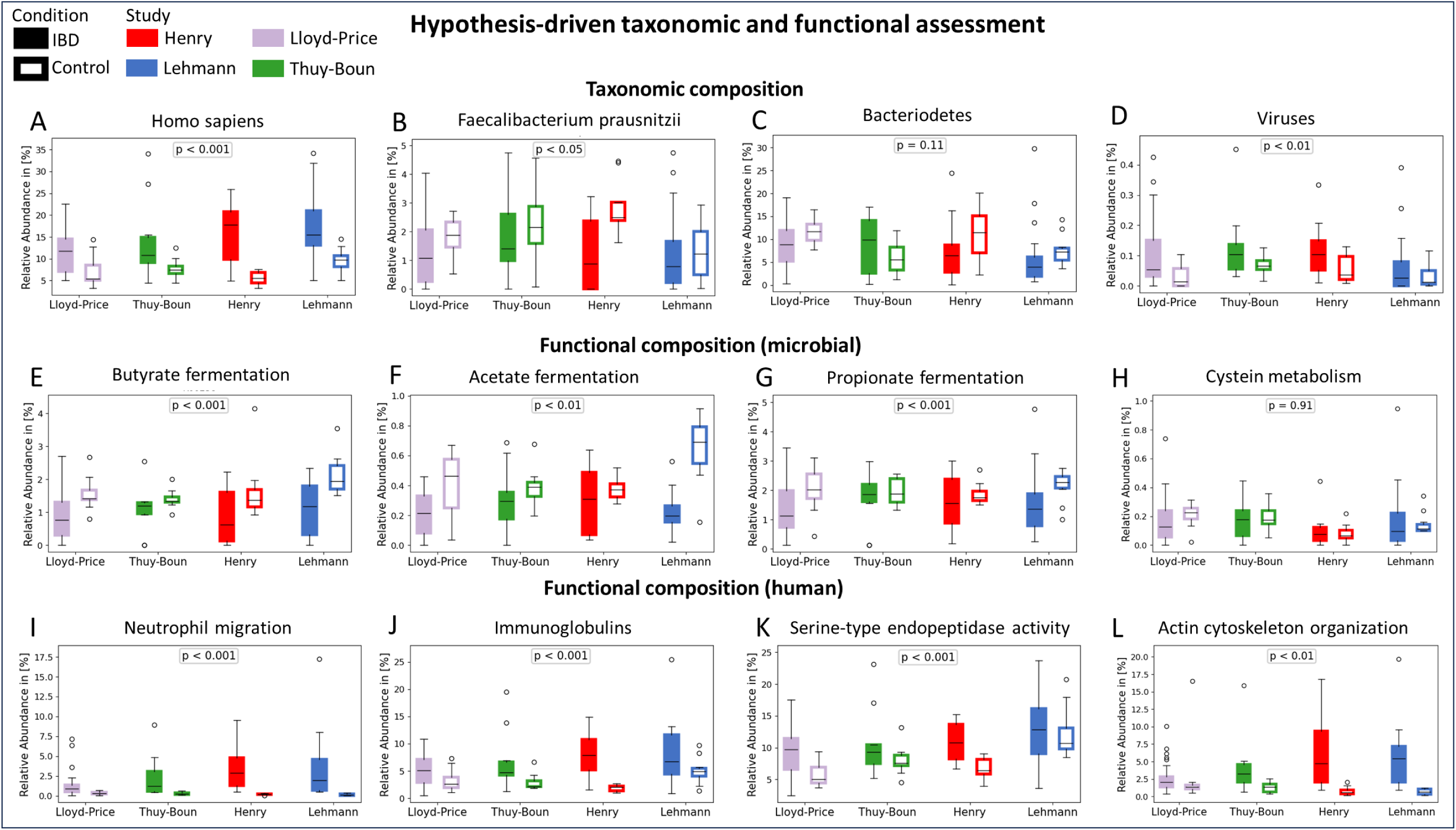
Hypothesis-driven taxonomic and functional assessment. Individual box-plots display the summed abundance of proteins grouped by their annotation (taxonomic annotation, functional annotation, incl. KEGG Reactions, GO:Terms or protein description). Colors imply the different studies. Filled boxes imply abundance in samples of IBD patients, while unfilled boxes represent the abundance in control samples. A – D: Abundance of the species *Homo sapiens* (A) and *Faecalibacteriumm prausnitzii* (B), the phylum *Bacteriodetes* (C) superkindom Virus (D). E – H: Abundance of microbial proteins involved in the reactions of butyrate (E), acetate (F) and propionate (G) fermentation as well as cysteine metabolism (H). The KEGG reactions included are specified in Supplementary File 6. I – L: Abundance of human proteins involved in neutrophil migration, immunoglobulins, serine-type endopeptidase activity and actin cytoskeleton organization. P-values were generated by mixed-effect models using Pythons statsmodels package.

One widely reported taxonomic shift in IBD is the reduced abundance of *Faecalibacterium prausnitzii*, a bacterial species considered to extert anti-inflammatory effects ^33^. Our meta-analysis confirms a statistically significant decrease in this species (p < 0.05, Figure 2B). Other proposed keystone taxa such as *Blautia*, *Ruminococcus*, and *Bifidobacterium* were also significantly decreased in IBD patients (Supplementary Figure 5-8). Another frequently discussed alteration is a change in the Firmicutes/Bacteroidetes ratio, although previous studies have reported conflicting results, several indicating an increase in Firmicutes and a decrease in Bacteroidetes, and others the opposite ^34–37^. Our analysis reveals a significant decrease in Firmicutes in IBD (Supplementary Figure 9). However, no consistent trend was observed for Bacteroidetes (p = 0.11, Figure 2C). These findings illustrate how meta-analyses can refine hypotheses and reduce the number of potential targets for interventions such as dietary modifications or therapeutic strategies.

Recent interest has emerged in antiviral therapies for IBD, supported by reports of increased viral abundance based, e.g., on RNA-Seq data ^38^. To our knowledge, an increase in viral abundance in IBD has not previously been reported using fecal metaproteomics. However, our results demonstrate a significant elevation of viral proteins in IBD samples (p < 0.01, Figure 2D), supporting the hypothesis of viral involvement in disease pathogenesis. Interestingly, we observed a significant increase in Caudovirales, a bacteriophage order, that was found associated with IBD in several studies ^32,39^ (Supplementary Figure 10).

Beyond taxonomic shifts, assessing functional composition is a key strength of metaproteomics. Fermentation of short-chain fatty acids (SCFAs), including butyrate, propionate, and acetate, is an essential anti-inflammatory process mediated by the gut microbiota ^40^. Consistent with prior reports ^41,42^, we observed a significant depletion of these fermentation pathways in IBD patients (Figure 2E-G). Conversely, although altered microbial cysteine metabolism has been reported in several studies, our data do not support this association (p = 0.91, Figure 2H). Since our analysis was performed at the whole-microbiome level and across all samples, taxon- or disease subtype-specific effects may be masked, as suggested by the findings of Zhang et al. ^32^ or Morgan et al. ^43^.

Key hallmarks of IBD, such as neutrophil infiltration and elevated immunoglobulin levels, were also reflected in our proteomic data. An increased abundance of human proteins involved in these immune processes was consistently observed across studies, resulting in a significant overall increase in the meta-analysis (Figure 2I-J). Moreover, we successfully validated a key finding from one of the included studies. Thuy-Boun et al. reported increased serine-type endopeptidase activity in IBD, which our meta-analysis confirmed as a generalizable feature across metaproteomic datasets (p < 0.001, Figure 2K). Additionally, we observed a significant increase of proteins involved in actin cytoskeleton organization in IBD patients (p < 0.01), previously identified as the most enriched process in the plasma proteome of IBD patients ^44^ (Figure 2L). This cross-validation between fecal and plasma proteomics suggests that cytoskeletal reorganization may serve as a central feature of gut inflammation.

### Functional and taxonomic annotation of potential biomarkers reveals identification of proteins connected to neutrophilic ROS response and IBD marker species

The set of 59 potential IBD biomarkers consisted of 24 human and 35 microbial metaproteins. All human proteins were increased in IBD patients, whereas all microbial metaproteins were higher abundant in control samples (Figure 3A). This finding aligns with the study of Thuy-Boun et al., which also found no host proteins in increased abundance in healthy subjects. An overall higher abundance of proteins of the immune system and a disrupted microbiome, leading to a higher relative abundance of host proteins, is often seen in IBD patients as confirmed in this meta-analysis.

**Figure 3.**
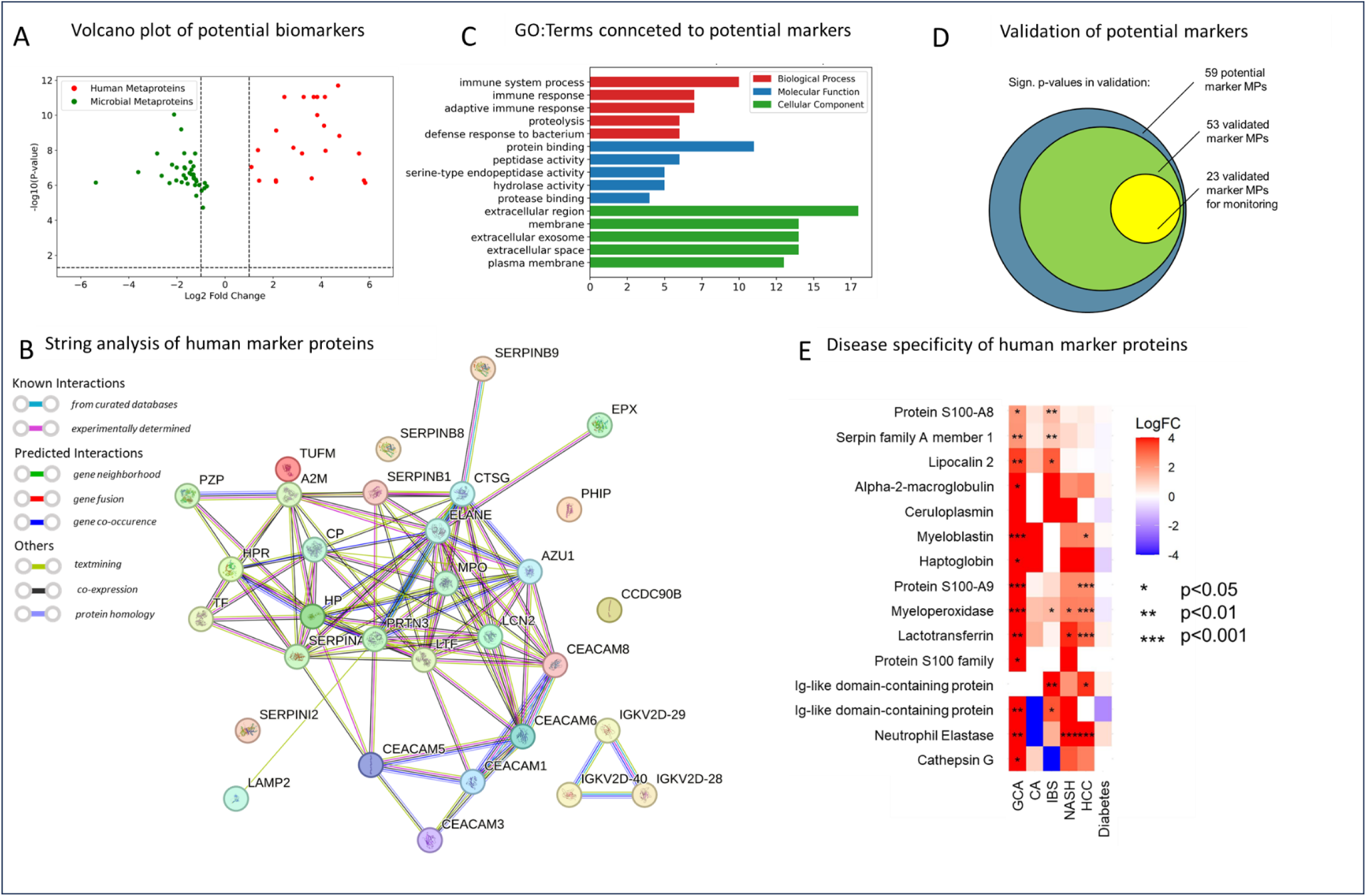
Biological interpretation, validation, and evaluation of disease specificity of potential biomarkers. A: Volcano plot comparison of potential marker metaproteins. B: String interaction network of human potential marker proteins. C: GO molecular function relative abundance plots. D: Venn plot of potential marker proteins that have significant p-values in one of the diagnosis studies (green) and that have significant p-values in remitting patients after therapy. E: Log fold change and p-value of potential human marker protein in non-IBD studies. GCA – Gastric carcinoma, CA – Colon adenoma, IBS – Irritable bowel syndrome, NASH – Non-alcoholic steatohepatitis, HCC – Hepatocell carcinoma.

#### Human Proteins

The identified altered host metaproteins were mainly neutrophil-derived, as shown by an extremely dense STRING network (Figure 3B), leading to a highly significant protein– protein interaction (PPI) enrichment (p-value: 0.0, 31 nodes, 99 edges, confidence score >400). In comparison to the STRING analysis performed in the study of Thuy-Boun (p-value: 1.84 × 10−12, 25 nodes, 26 edges, confidence score >400), the meta-analysis approach increased the confidence in the functional associations of the potential biomarkers.

Many of the human proteins identified in this meta-analysis are components of neutrophil-derived extracellular vesicles and participate in essential hallmarks of IBD immune response ^45^. For example, Ceruloplasmin, Lipocalin-2 and the Calprotectin subunits Protein S100-A8/A9 are involved in the sequestration of iron, thereby preventing on the one hand Fenton reaction and production of reactive oxygen species, and on the other hand withholding an essential element for microbial growth.

The highest enriched pathways resulting from the STRING analysis were “defense responses against bacterium” and “neutrophil response”, which is in line with the metaproteomic analysis of extracellular vesicles in IBD patients conducted by Zhang et al. and the results of the study of Thuy-Boun ^13,32^.

Only a proportion of these biomarkers were explicitly described in the four original studies. Whereas the neutrophil-derived human proteins S100-A8/A9, Myeloperoxidase, and Azurocidin were described in three out of four studies, we were able to validate the increased abundance of several other proteins, only mentioned in one or two of the studies, including Cathepsin (Table 1). Other potential markers found in the meta-analysis were not described in the underlying metaproteomic studies, but were found increased, e.g., in serum of IBD patients, like Alpha-1-acid glycoprotein ^46^. This protein is a major acute phase protein, whose abundance increases in response to tissue injury or inflammation ^47^.

**Table 1.**
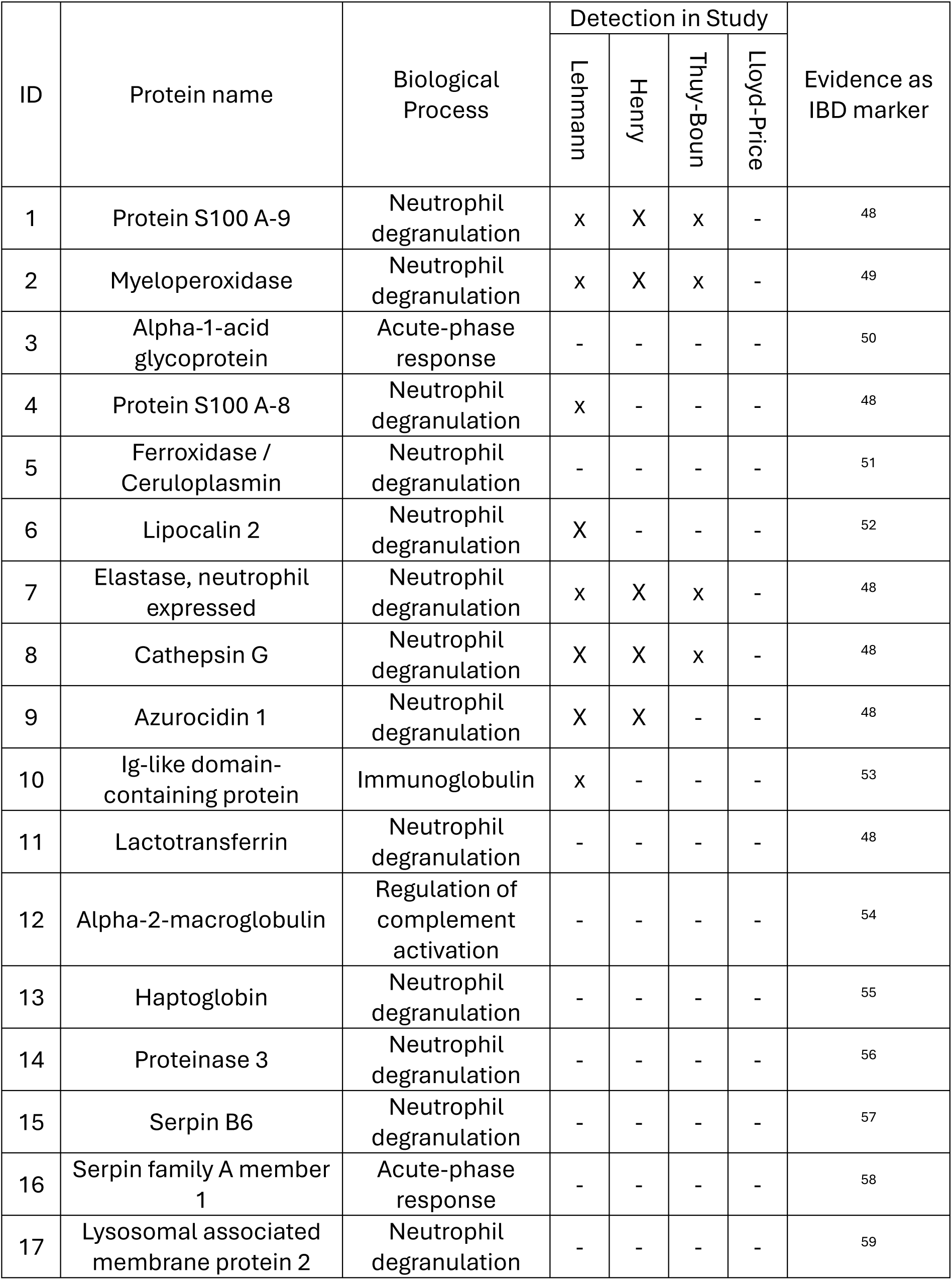
Reporting of potential human marker proteins in the discovery studies. The protein name is derived from Unipept-Annotation or Uniprot-IDs in metaproteins. The detection in study describes, whether the proteins were mentioned in the published manuscript of the dataset. Metaproteins 37164 was excluded due to conflicting annotations from Prophane and Unipept. Immunoglobulin-domain containing proteins were merged into one entry.

| ID | Protein name | Biological Process | Detection in Study |  |  |  | Evidence as IBD marker |
| --- | --- | --- | --- | --- | --- | --- | --- |
|  |  |  | Lehmann | Henry | Thuy-Boun | Lloyd-Price |  |
| 1 | Protein S100 A-9 | Neutrophil degranulation | x | X | x | - | 48 |
| 2 | Myeloperoxidase | Neutrophil degranulation | x | X | x | - | 49 |
| 3 | Alpha-1-acid glycoprotein | Acute-phase response | - | - | - | - | 50 |
| 4 | Protein S100 A-8 | Neutrophil degranulation | x | - | - | - | 48 |
| 5 | Ferroxidase / Ceruloplasmin | Neutrophil degranulation | - | - | - | - | 51 |
| 6 | Lipocalin 2 | Neutrophil degranulation | X | - | - | - | 52 |
| 7 | Elastase, neutrophil expressed | Neutrophil degranulation | x | X | x | - | 48 |
| 8 | Cathepsin G | Neutrophil degranulation | X | X | x | - | 48 |
| 9 | Azurocidin 1 | Neutrophil degranulation | X | X | - | - | 48 |
| 10 | Ig-like domain-containing protein | Immunoglobulin | x | - | - | - | 53 |
| 11 | Lactotransferrin | Neutrophil degranulation | - | - | - | - | 48 |
| 12 | Alpha-2-macroglobulin | Regulation of complement activation | - | - | - | - | 54 |
| 13 | Haptoglobin | Neutrophil degranulation | - | - | - | - | 55 |
| 14 | Proteinase 3 | Neutrophil degranulation | - | - | - | - | 56 |
| 15 | Serpin B6 | Neutrophil degranulation | - | - | - | - | 57 |
| 16 | Serpin family A member 1 | Acute-phase response | - | - | - | - | 58 |
| 17 | Lysosomal associated membrane protein 2 | Neutrophil degranulation | - | - | - | - | 59 |

A level of evidence for the potential involvement of the protein in IBD is included in Table 1.

The GO:Terms connected to the human proteins were “immune system process” (Biological Process), “protein binding” (Molecular function”) and “extracellular region” (Cellular Component, Figure 3C). In general, the connected GO:Terms portray key aspects of IBD pathogenesis and confirm findings, that were described in multiple publications before. For example, the importance of extracellular vesicles and exosomes was previously evaluated by Zhang et al. and protein-binding is crucial for immune reactions, e.g., antibody-binding or acute-phase or complement proteins ^32^. Furthermore, proteolysis is explicitly described in IBD pathogenesis ^60^.

Apart from GO:Terms the graph database enables the analysis of drugs connected to the potential human marker proteins. Here, five proteins were connected to Tretinoin (Supplementary File 7), also known as all-trans retinoic acid, which was described to attenuate inflammation in colonic inflammation in several publications ^61,62^.

#### Microbial Proteins

The great majority of potential microbial marker proteins were annotated to short-chain fatty acid (SCFA) producers, e.g., *Lachnospiraceae* and *Clostridiaceae*, genera *Blautia*, *Roseburia*, and *Ruminococcus*. SCFAs, most importantly butyrate, are known to have anti-inflammatory properties, e.g., through the inhibition of proinflammatory cytokines ^63^. Most prominently, three proteins were attributed to *Faecalibacteria* species, which are described as decreased in IBD in numerous studies ^33,64^. Especially, proteins related to oxidative stress response and iron-/ metal-binding potential, like rubrerythrin, NADH peroxidase or Iron-containing alcohol dehydrogenase were decreased in IBD patients. As many human proteins that showed increased abundance in IBD patients, are involved in iron-sequestration and oxidative stress, this interaction is identified as a key process by this meta-analysis. Downregulation of specific genes, including rubrerythrin was observed in gut microbial species before, and is part of a modulation of their metabolism promoting resilience during inflammation-induced iron deficiency ^65^. While iron supplementation-based therapies are evaluated in IBD, the interactions of these therapies with the microbiome are complex, potentially even exacerbating the inflammation ^66^

### Potential biomarkers partially validated

Many metaproteomic studies propose potential biomarkers but lack validation with independent datasets. The meta-analysis approach bears potential to validate biomarkers in independent samples. Here, a subset of samples from the Lloyd-Price dataset and an in-house dataset comprising IBD patients and controls was used for validation by testing for significantly altered abundance of these metaproteins. Furthermore, the application of the metaproteins as a marker for therapy monitoring was assessed by testing abundance in a dataset of IBD patients before and after the therapy (n=24).

53 of 59 metaproteins were found to be significantly changed in at least one of the validation data sets (Figure 3D). Ten metaproteins, including Alpha-1-acid glycoprotein, an immunoglobulin, and several *Lachnospiraceae* metaproteins were significantly differentially abundant in both datasets. Alpha-1-acid glycoprotein was once tested as a plasma biomarker, but achieved only limited sensitivity ^46^. Furthermore, recent studies ^67^ support a relationship between IBD and *Lachnospiraceae*, which were also reported to be associated with lower butyric acid levels in IBD ^68^. Our results suggest its usefulness for IBD diagnosis in fecal samples. Furthermore, 23 of the 53 validated metaproteins were significantly changed in IBD patients achieving remission after biologics therapy. A large proportion is associated with neutrophil degranulation.

### Marker proteins not specific for IBD

Alongside validation, exploring the disease specificity is an important aspect of biomarker discovery. IBD in particular shares several symptoms with other diseases, like Irritable Bowel Syndrome (IBS), or diabetes. While Calprotectin is widely used in clinical settings, its use is not optimal, as it was elevated in several other conditions, like viral or bacterial infections ^69,70^, necrotizing enterocolitis ^71^ or graft-versus-host disease ^72^. Furthermore, IBD can develop into or propagate the development of gastrointestinal cancers. A differentiation between other disorders and IBD is of tremendous importance, e.g., for the development of therapy approaches. Therefore, we inspected the abundance of the potential biomarkers in other diseases like IBS, NASH, HCC, CA, and diabetes. In Figure 3E, fold changes of a subset of the human potential marker proteins are depicted. (The microbial proteins are depicted in Supplementary Figure 11). In several diseases a large subset of the potential marker metaproteins significantly changed as well. For instance, Neutrophil Elastase (NE) is significantly higher abundant in, e.g., gastric cancer, NASH, and HCC. NE was described as important for NASH development ^73^ and it can facilitate tumor growth ^74^, therefore the results of our analysis are at least partially confirmed. On the other hand, the data implies that NE is lower abundant in CA, whereas Cathepsin G is lower abundant in IBS. As NE was reported as a biomarker of colon cancer ^75^, this trend might be only suitable for this specific patient set, and larger clinical cohorts are needed for a general statement of disease specificity. Conversely, Cathepsin G, which is released by neutrophils, could be used for discrimination between IBD and IBS, as it could be seen as a marker for neutrophil infiltration, which is an efficient differentiator between both pathologies. In diabetes, most metaproteins show only minor changes.

Therefore, a subset of these proteins could be used to exclude certain conditions. This supports the need for complex biomarker panels rather than single proteins to achieve a differential diagnosis.

### Metaproteomics meta-analysis reveals common and differentiating features between UC and CD

An important goal of recent IBD research is to find specific biomarkers to differentiate between CD and UC ^76^, as they share symptoms, genotypes and expression profiles ^77^, but an exact diagnosis is central for successful therapy selection. Analysis of variance sources revealed that 23 microbial and 19 non-microbial metaproteins exhibit differential abundance in comparison to healthy controls, associated with both UC and CD, respectively (Figure 4A-D). In contrast, several metaproteins displayed CD or UC-specific variance. In CD, 27 microbial and five non-microbial metaproteins showed a proportion of variance explained by disease greater than 0.2, whereas in UC, 17 microbial and 38 non-microbial metaproteins met this threshold. Notably, the UC-associated set included a substantial number of neutrophil-derived proteins. The higher number of microbial metaproteins differentially abundant due to the disease in CD hints at a more pronounced dysbiosis in CD compared to UC, which was observed in several other publications as well ^78,79^. In contrast, the number of infiltrating neutrophils and differentially changed immune proteins and NET formation is considered to be higher in UC ^80,81^, which would explain its stronger association with human proteins. While there are more microbial metaproteins associated with CD, the most prevalent microbial family in both sets unique for the disease subtype was *Lachnospiraceae*. All metaproteins of *Lachnospiraceae* taxa were decreased in abundance in both subtypes. Decreased abundances of *Lachnospiraceae* was observed in active CD as well as UC and connected to a decreased butyrogenesis ^68,82,83^. Furthermore, there was no bias to a specific pathway or enzyme class in both metaprotein sets (Supplementary File 8). These results indicate a high concordance in the metaproteomic profiles associated with the two IBD subtypes, with differences primarily reflecting the relative severity of disease manifestations between them.

**Figure 4.**
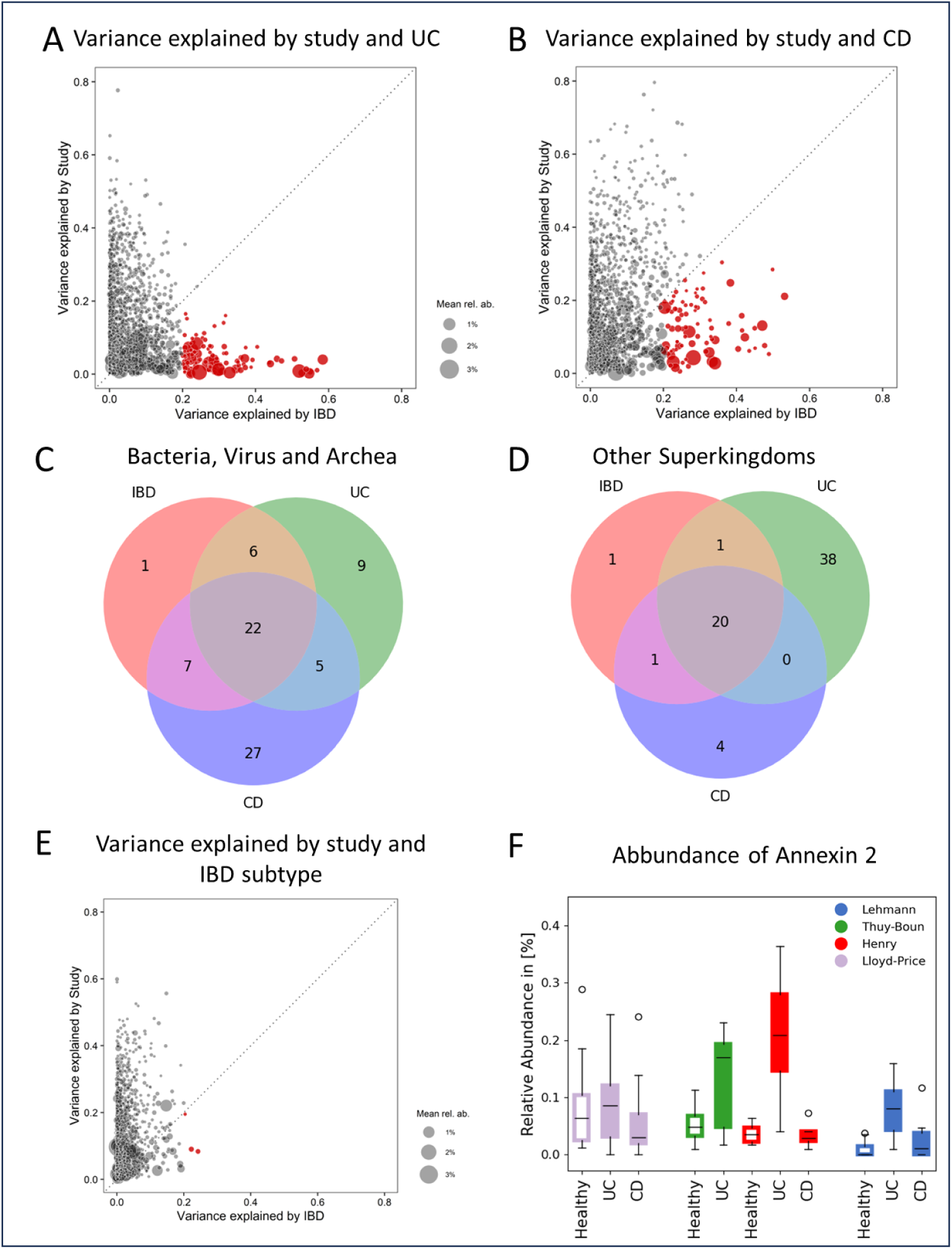
Meta-analysis of IBD subtype specificity of microbial and non-microbial metaproteins. A: A: Variance explained by the study compared to variance explained by UC for all metaproteins identified in all studies. 101 metaproteins with variance explained by UC > 0.2 were marked red. B: Variance explained by the study compared to variance explained by CD for all metaproteins identified in all studies. 86 metaproteins with variance explained by CD > 0.2 were marked red. C: Venn diagram representing shared and unique microbial (viral, bacterial, archeal) metaproteins with variance explained by UC, CD and IBD > 0.2. D: Venn diagram representing shared and unique non-microbial metaproteins with variance explained by UC, CD and IBD > 0.2. E: Variance explained by the study compared to variance explained by UC for all metaproteins identified in all studies. Three metaproteins with variance explained by UC compared to CD > 0.2 were marked red. F: Abundance of human Annexin 2 (Meta-Protein 8524) in different disease subtypes and studies.

The most promising metaprotein for differentiation between UC and CD to report is Annexin 2, which has the highest variance explained by disease subtype (Figure 4E). Annexin 2 regulates intestinal epithelial cell spreading and wound healing and was described as differentiating factor by Zhang et al. ^84,85^. In line with the findings of this paper, Annexin 2 was upregulated in UC in comparison to CD patients and healthy controls in three of the discovery cohorts (Figure 4F). In recent evaluations of fecal biomarkers for differentiation of CD and UC, Annexin 2 was not considered ^76,86^. Two other metaproteins associated with the disease subtypes were Fibrinogen alpha chain and Fibrinogen beta chain, which were both higher abundant in UC than in CD and healthy individuals. Conversely, Fibrinogen was described elevated in active disease in both, UC and CD ^87^.

## Discussion

This meta-analysis identified numerous potential human and microbial diagnostic protein biomarkers for inflammatory bowel diseases (IBD). To enable this, raw data from ten publicly available and in-house metaproteomic datasets were reanalyzed and integrated using a unified bioinformatic workflow.

Most of the identified human proteins were linked to neutrophilic granulocytes, which have been described as the main drivers of inflammation in IBD ^88^. The microbial proteins, on the other hand, were primarily involved in defense against reactive oxygen species and associated with microbial taxa known to be linked to inflammation. In an additional step, most metaproteins extracted in the discovery cohorts were confirmed in at least one of two validation cohorts. The analysis of disease specificity implies the need for larger panels of biomarkers instead of individual ones to differentiate IBD from other disorders.

The used approach demonstrated the value of data reuse from public repositories for biomarker discovery, validation, and assessing disease specificity. It highlights the importance of meta-analysis in microbiome research, as this approach helps in narrowing and prioritizing hypotheses without requiring significant additional measurement effort or investment ^31^. Both, well-established biomarkers (such as LTF and S100-A8/A9) and less-explored or even novel human proteins, were identified as differentially expressed in IBD or differentiating UC and CD, highlighting their potential for diagnostics and disease monitoring. However, identifying generalizable protein biomarkers using mass spectrometry remains a significant challenge. One of the main issues is the limited overlap of metaproteins identified across studies. This issue can be attributed to confounding factors such as differing experimental workflows, like those evaluated in CAMPI studies ^29^, and interindividual variation, as well as the lack of standardization in metaproteomic workflows and sample preservation (CAMPI02), which remains in its early stages. The emergence of data-independent acquisition (DIA) methods in metaproteomics offers a promising path forward ^89^. Unlike Top-n data-dependent acquisition (DDA) approaches, which may miss many peptides in complex stool samples, DIA could significantly reduce the sparseness of data in future meta-analyses. This strategy may lead to the discovery of more potential biomarker metaproteins. Another persistent challenge is the use of large sequence databases and varying protein grouping strategies, which can influence identification outcomes ^90^. Additionally, comprehensive and standardized metadata annotation is essential for the reanalysis of repository data, a need that the metaproteomics community has recently begun to address. Although (meta-)proteomic meta-analyses remain relatively scarce compared to the more common (meta-)genomic ones, they are beginning to gain attention. This growing interest underscores the importance and potential of metaproteomics in understanding disease mechanisms and developing clinically relevant biomarkers for conditions like IBD. The extension of this meta-analysis to other sample types (e.g., saliva, biopsies) might be a promising path to explore human-microbiome interactions in other diseases.

## Conclusion

Taking together, we were able to integrate and analyze several fecal metaproteomic datasets. After the extraction of metaproteins whose abundance was largely affected by IBD, not by study-specific effects, iron sequestration and oxidative stress response were confirmed as essential human microbiome interactions in IBD. We were able to confirm several biomarkers reported before, but also discovered new fecal markers, like Alpha-1-acid glycoprotein. Furthermore, we validated the majority of biomarkers and showed that biomarker panels are most likely needed to distinguish IBD from other disorders.

## Acknowledgments

The Inflammatory Bowel Disease Family Cohort (IBD-FC) received funding by the German Research Foundation (DFG) through Excellence Clusters “Inflammation at interfaces” (EXC 306) and “Precision Medicine in Inflammation” (PMI; EXC 2167), as well as of the Research Unit miTarget (FOR 5042). It was also supported by a grant from the German Ministry of Education and Research (01ZX1606A). K.A. received funding from the EKFS research grant #2019_A09 and EKFS Clinician Scientist Professorship (K.A., 2020_EKCS.11), the BMBF (eMED Juniorverbund “Try-IBD” 01ZX1915A, 01ZX2215, K.A.), the DFG RU5042 (K.A.), the Joachim Herz Stiftung (K.A.).

## Authors’ contributions

M.W. and R.H. coordinated development and construction of the manuscript, with input from all co-authors. R.H. supervised this study. M.W., L.K., K.S., A.G. and P.H. were involved in data processing and bioinformatics analyses. All co-authors contributed edits and input throughout the manuscript construction, and all have read and approved the final version.

## Abbreviations

ANOVA: Analysis of Variance
API: Application Programming Interface
CA: Colon Adenoma
DDA: Data-dependent aquisition
DIA: Data-independent aquisition
E.C.: Enzyme Commission
GCA: Gastric Carcinoma
GO: Gene Ontology
HCC: Hepatocellular Carcinoma
IBD: Inflammatory Bowel Disease
IBS: Irritable Bowel Syndrome
KEGG: Kyoto Encyclopedia of Genes and Genomes
LCA: Lowest Common Ancestor
NASH: Nonalcoholic Steatohepatitis
NCBI: National Center for Biotechnology Information
NE: Neutrophil Elastase
Permanova: Permutational Analysis of Variance
PPI: Protein-Protein-Interaction Network
SCFA: Short-Chain Fatty Acid

