## Supplementary Figures for "A metaproteomics-based meta study of samples from patients with inflammatory bowel disease identifies potential markers for diagnosis and therapy monitoring"

Wolf et al.

Supplementary Information

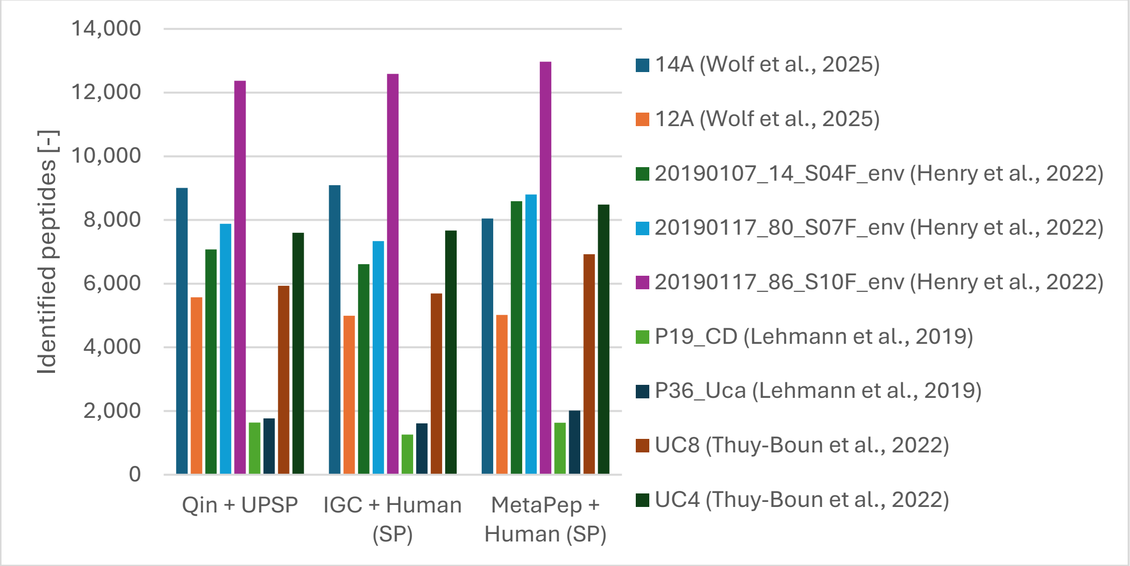

*Supplementary Figure 1 Number of identified peptides using different sequence databases at 1% False discovery rate from a set of samples from different studies.* The used databases are 1) a metagenome catalogue published by Qin et al. (DOI: 10.1038/nature08821) concatenated with the Uniprot/SwissProt database, 2) a metagenome catalogue published by Li et al. (DOI: 10.1038/nbt.2942) concatenated with human SwissProt entries, and 3) a microbial peptide catalogue published by Sun et al. (DOI: 10.1016/j.csbj.2023.08.025) concatenated with human SwissProt entries.

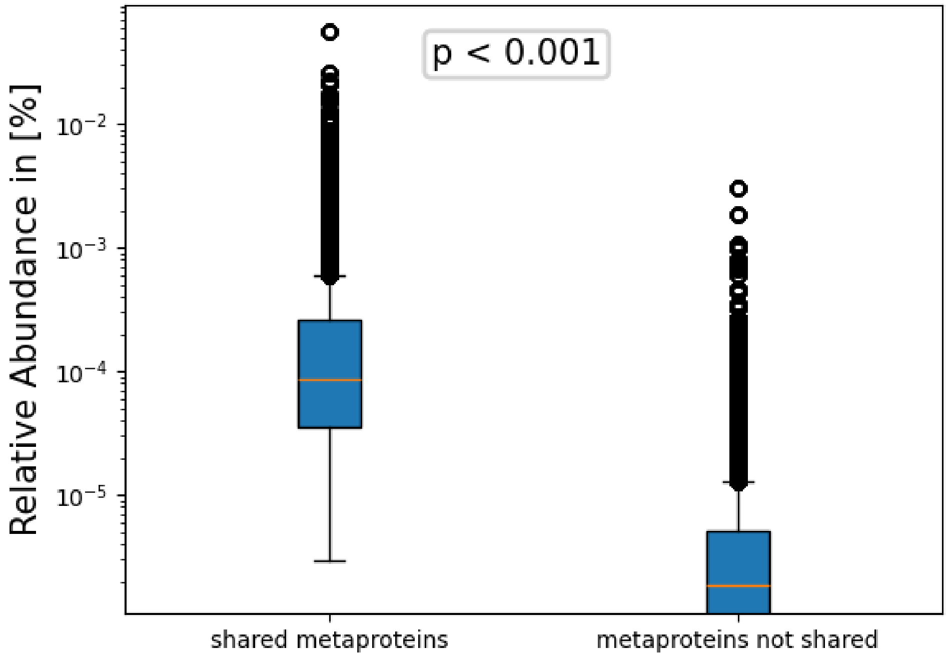

*Supplementary Figure 2 Average abundance of metaproteins identified in all discovery datasets ("shared metaproteins") and metaproteins that were not identified in all discovery datasets ("metaproteins not shared"). P-value was calculated using Mann-Whitney U test.*

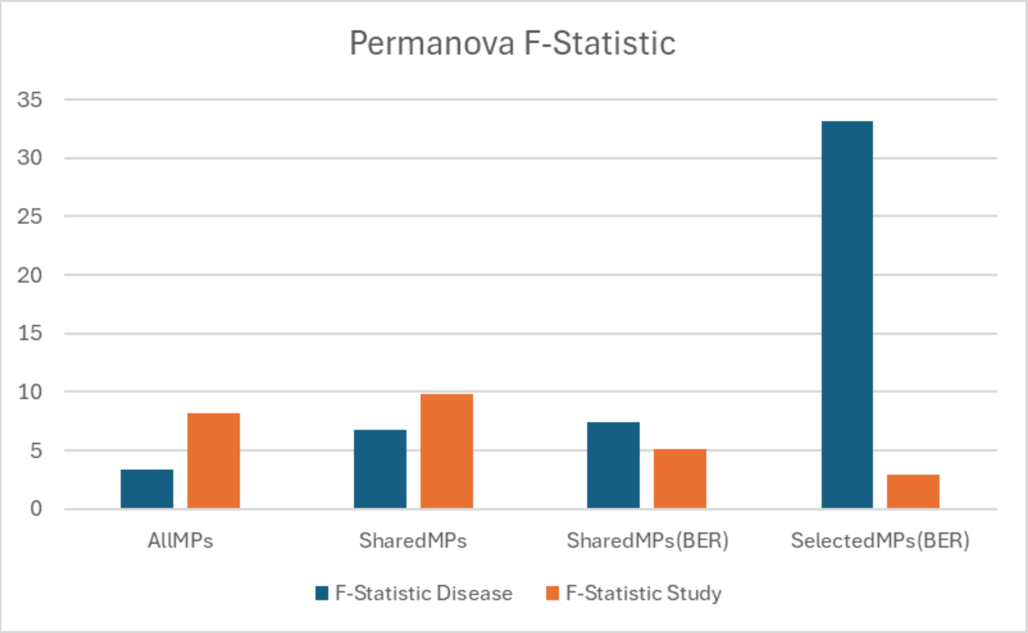

*Supplementary Figure 3 PERMANOVA pseudo-F-statistics, corresponding to importance of the grouping factor (healthy/IBD and differnte studies), using all identified metaproteins, all metaproteins identified in each study of the discovery dataset, all metaproteins identified in each study of the discovery dataset after the batch effect reduction (BER) and* all 59 metaproteins with variance explained by IBD > 0.2 after batch effect reduction.

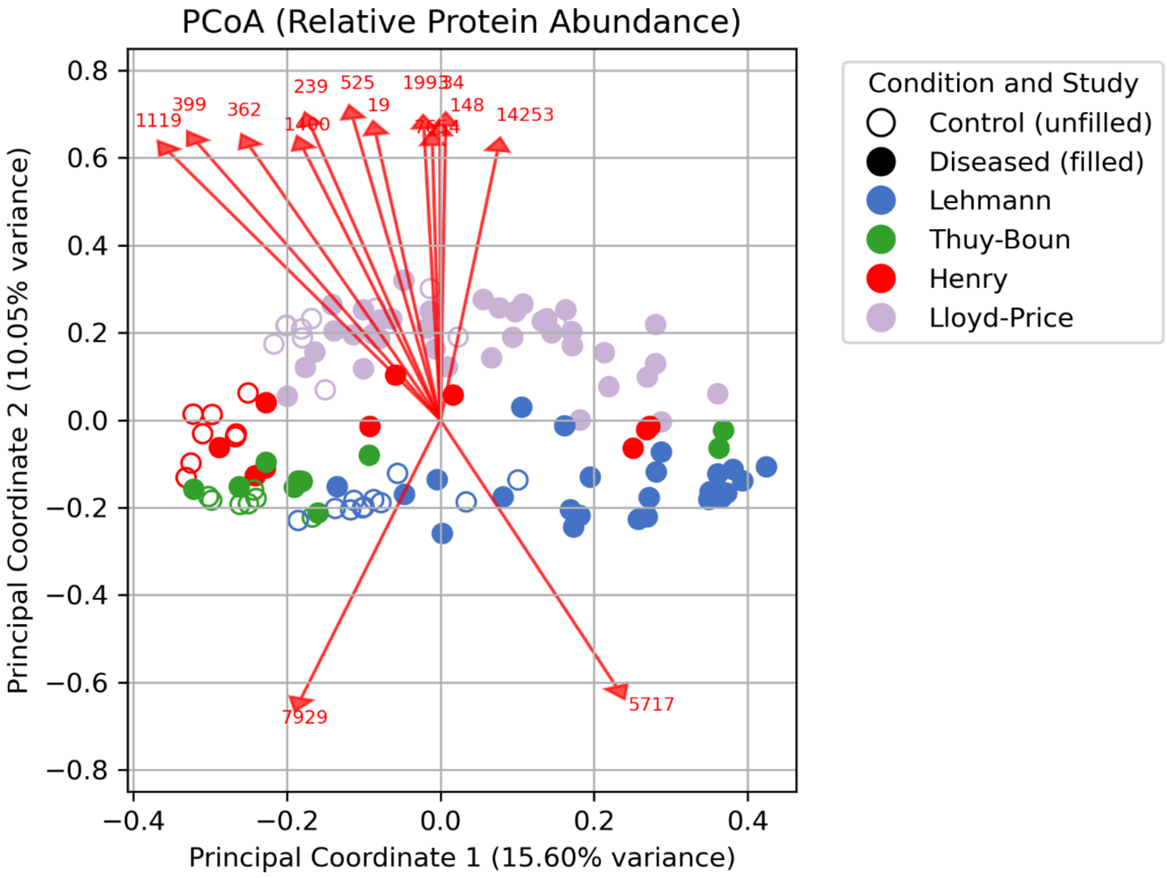

*Supplementary Figure 4 : Principle Coordinate Analysis (PcoA) score plot of metaproteins identified in all studies from the discovery cohort*. Arrows represent the metaproteins with the strongest correlation with Principle Coordinate 2.

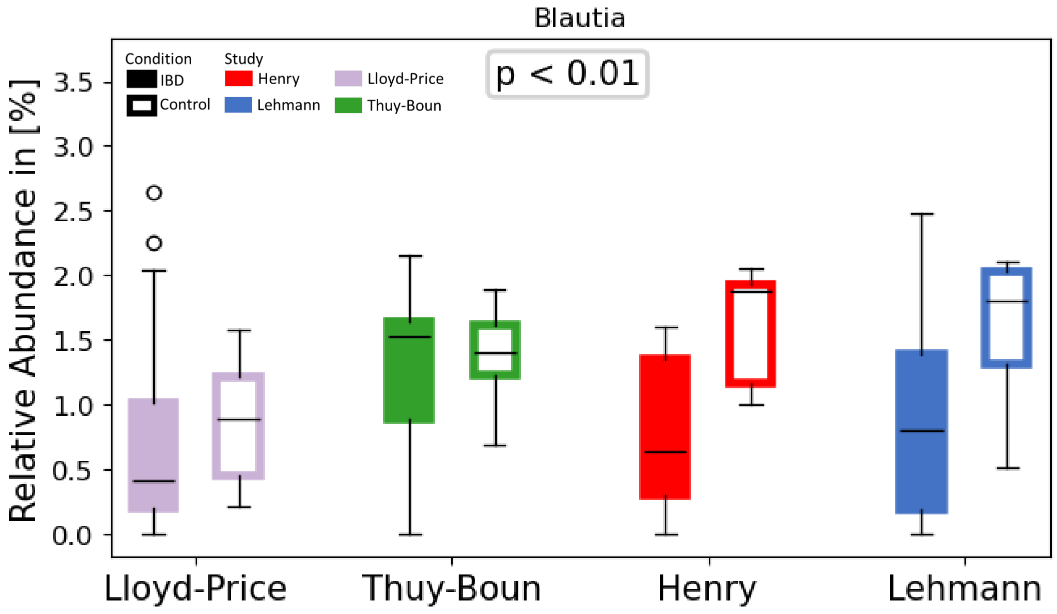

*Supplementary Figure 5 Abundance of metaproteins the genus Blautia*. Colors imply the different studies. Filled boxes imply abundance in samples of IBD patients, while unfilled boxes represent the abundance in control samples.

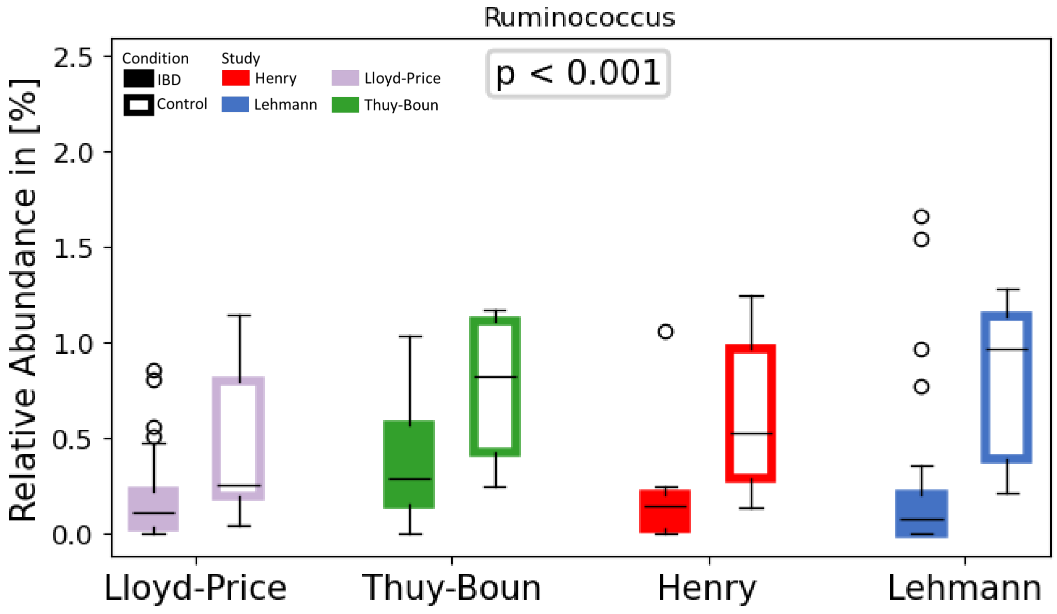

*Supplementary Figure 6 Abundance of metaproteins the genus Ruminococcus*. Colors imply the different studies. Filled boxes imply abundance in samples of IBD patients, while unfilled boxes represent the abundance in control samples.

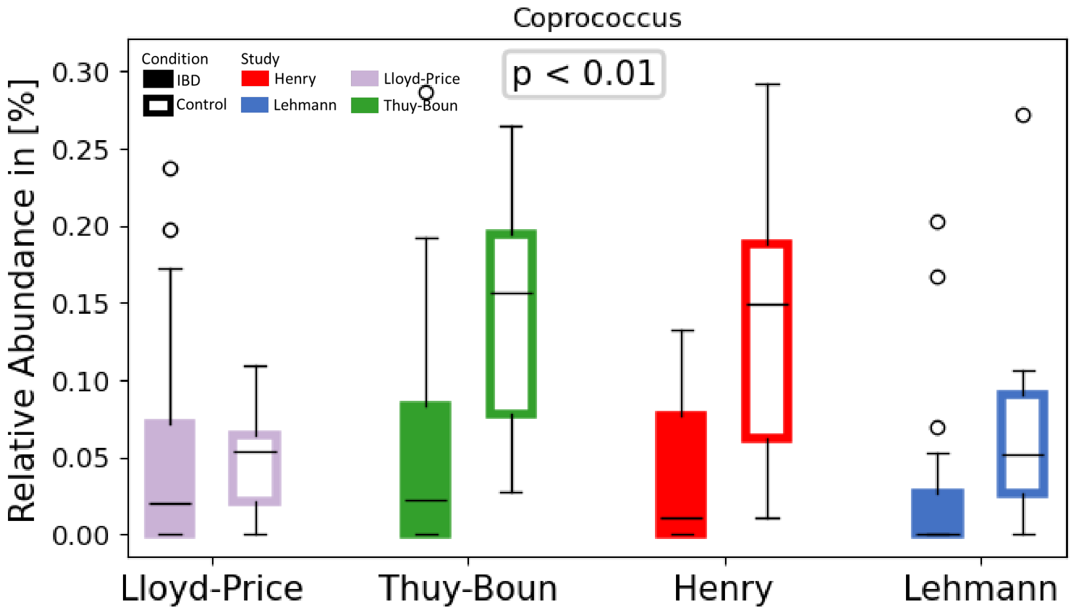

*Supplementary Figure 7 Abundance of metaproteins the genus Coprococcus*. Colors imply the different studies. Filled boxes imply abundance in samples of IBD patients, while unfilled boxes represent the abundance in control samples.

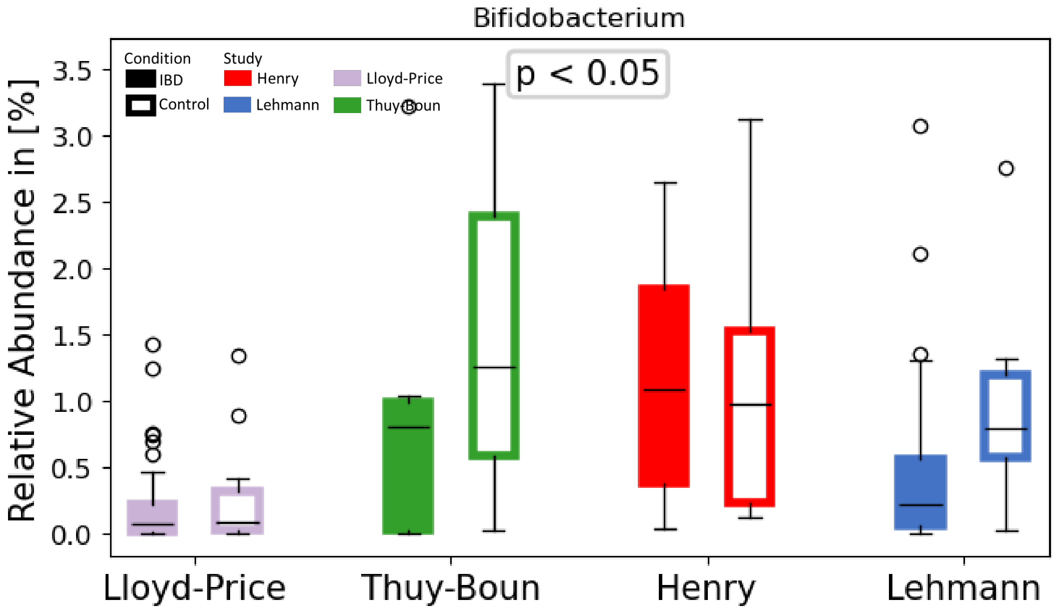

*Supplementary Figure 8 Abundance of metaproteins the genus Bifidobacterium*. Colors imply the different studies. Filled boxes imply abundance in samples of IBD patients, while unfilled boxes represent the abundance in control samples.

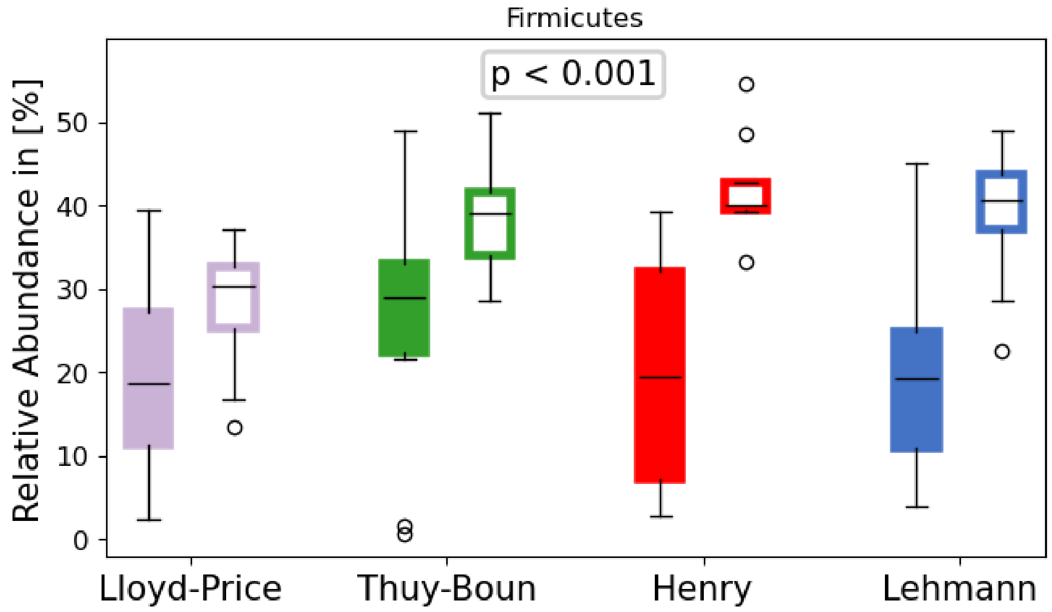

*Supplementary Figure 9 Abundance of metaproteins the phylum Firmicutes.* Colors imply the different studies. Filled boxes imply abundance in samples of IBD patients, while unfilled boxes represent the abundance in control samples.

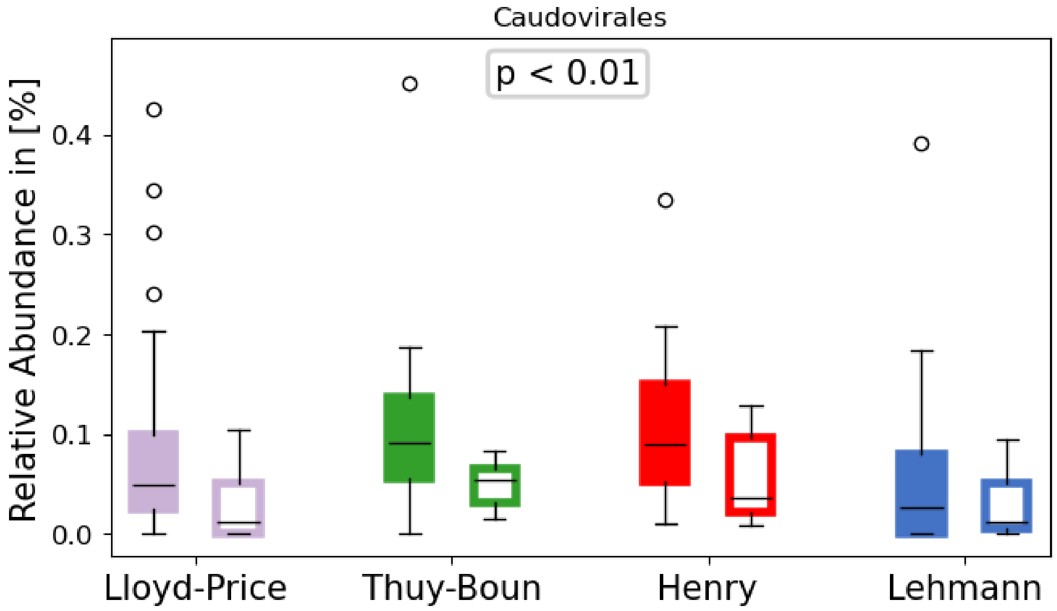

*Supplementary Figure 10 Abundance of metaproteins the class Caudovirales.* Colors imply the different studies. Filled boxes imply abundance in samples of IBD patients, while unfilled boxes represent the abundance in control samples.

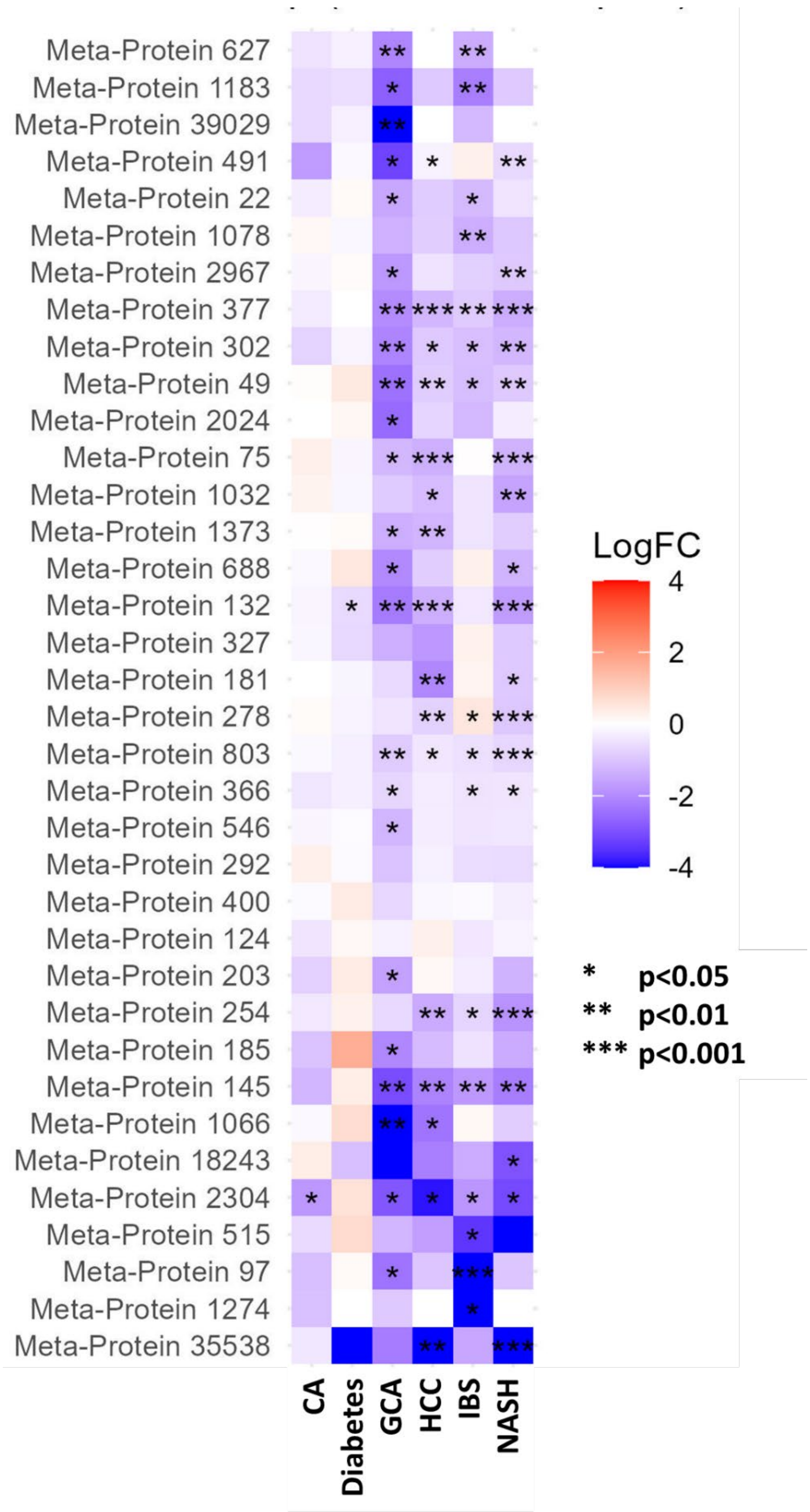

*Supplementary Figure 11 Heatmap (Log fold change and p-value) of potential microbial marker protein in non-IBD studies.* GCA – Gastric carcinoma, CA – Colon adenoma, IBS – Irritable bowel syndrome, NASH – Non-alcoholic steatohepatitis, HCC – Hepatocell carcinoma. *P-value were calculated using Mann-Whitney U test.*
